# Associations Between Habitual Light Exposure-Related Behaviors and Sleep Timing and Sleep Complaints in an International Community Sample

**DOI:** 10.64898/2026.01.02.697416

**Authors:** Ann-Sophie Loock, Rafael Lazar, Manuel Spitschan, Christine Blume

## Abstract

Sleep is essential for health and light is an important environmental signal influencing its timing, quality, and regulation. Retinal light exposure reflects the interplay between environmental illumination and behavioral choices, yet it remains unclear which habitual light exposure-related behaviors meaningfully impact sleep outcomes. In this preregistered analysis, we examined associations between these behaviors, sleep timing, and sleep complaints in a large, international community sample (*N* = 774, *M*_age_ = 32.6 ± 14.6 years). Participants completed the Light Exposure Behavior Assessment (LEBA), with four behavioral domains included in the analyses. Sleep timing, sleep disturbances and sleep-related daytime impairment were measured using established questionnaires. Bayesian analyses indicated that time spent outdoors and device use in bed were most strongly associated with sleep outcomes. Greater time outdoors was linked to earlier sleep timing and fewer sleep complaints, whereas more frequent device use in bed was associated with greater sleep disturbance and daytime impairment. Morning and daytime lighting practices and evening light control showed no conclusive evidence. Together, these findings highlight the relevance of everyday light exposure-related behaviors for sleep and support behavioral approaches to promoting healthy sleep in real-world contexts.

## Introduction

Insufficient or disturbed sleep impairs attention, learning, and emotion regulation, and even moderate chronic reductions in sleep quantity or quality can affect daytime functioning or contribute to long-term health deterioration^1–3^. Sleep complaints are common, affecting roughly one-third to one-half of adults worldwide^4–6^. At both the individual and societal level, sleep problems contribute to physical and psychological health problems, reduced productivity, and higher healthcare costs^7–9^. Given these consequences, it is essential to identify environmental factors that shape sleep timing and sleep quality, and to understand how everyday behaviors influence exposure to them.

Among the environmental factors affecting sleep, light is one of the most potent and well-characterized regulators. Our sleep-wake pattern reflects the interaction of a circadian process, which aligns sleep propensity with internal biological time, and a homeostatic process tracking prior wakefulness and sleep need^10^. Light synchronizes the circadian system to the 24-hour day via intrinsically photosensitive retinal ganglion cells (ipRGCs) projecting to the suprachiasmatic nucleus, thereby influencing when sleep is biologically favored and when wakefulness is promoted^11–13^. Beyond its entraining role, light also exerts acute, non-circadian effects on arousal and sleep propensity through melanopsin-dependent pathways, modulating sleep latency, continuity, and subjective sleepiness^14–16^. The impact of light on sleep therefore depends strongly on its timing and intensity: morning and daytime light increases alertness, reduces sleepiness, supports consolidated wakefulness, and stabilizes downstream sleep^17,18^. The quality and architecture of sleep is also related to preceding light exposure with early light exposure to high intensities being associated with beneficial outcomes for deeper and better quality sleep at night^19,20^. In contrast, evening or nighttime light, particularly short-wavelength or bright artificial light, directly activates alertness pathways and suppresses melatonin, which has been suggested to lead to delayed sleep onset and increased sleep fragmentation^20^. The direct alerting effect of light in the evening can also systematically shorten sleep duration due to delayed sleep timing and fixed wake requirements^21^. Conversely, reducing light before bedtime can advance melatonin onset and improve sleep efficiency^22^. These circadian and direct alerting effects together make everyday light exposure patterns a central determinant of sleep timing and the experience of sleep disturbances and daytime functioning.

In modern daily life, light exposure is no longer dictated solely by the natural light dark cycle but is strongly shaped by individual behavior. People spend comparatively little time outdoors, on average around 1.5h on workdays^23^, which results in reduced daytime light exposure and, in turn, substantial artificial light in the evening^24–26^. This shift towards dimmer days and brighter evenings creates exposure patterns that diverge from those under which human circadian and systems evolved, and has been associated with later sleep timing and more sleep disturbances^27–29^. Daytime light exposure can also modulate sensitivity to evening light, with higher prior light exposure reducing the magnitude of melatonin suppression at night^30^. Although evening artificial light reliably suppresses melatonin, its downstream effects on sleep appear less pronounced than once assumed, as melatonin levels typically recover quickly once the light stimulus is removed and alterations in sleep timing or structure have been small or absent^31,32^.

Because modern light exposure patterns arise from both environmental illumination and individual choices, recent conceptual frameworks increasingly emphasize that light exposure should be understood as a behavior in its own right^33^. This perspective recognizes that physiological effects of light depend not only on the external light environment but also on how individuals interact with it and, finally, on the irradiance reaching the retina. Retinal irradiance is shaped by factors such as gaze direction, pupil size, and ocular filtering^34^, and by everyday behaviors including time spent outdoors, screen use, and the management of light sources. Despite its relevance, these behavioral patterns have rarely been examined systematically. Self-report instruments therefore offer a practical way to capture habitual, context-dependent light exposure-related behaviors at scale. Recently, a validated questionnaire comprehensively assessing such behaviors became available: the Light Exposure Behavior Assessment (LEBA) questionnaire^35^, which distinguishes five domains of behavior shaping light exposure.

The LEBA has so far been used in one empirical study linking its behavioral domains to chronotype and global sleep quality in a Malaysian sample^36^, and more recently in work combining self-report with wearable light sensors to examine how these behaviors relate to objectively measured light exposure^37,38^. However, no study has yet examined LEBA-derived behaviors in a large, international sample with respect to specific sleep outcomes such as sleep timing or night-time and daytime complaints. The present study addresses this gap by applying the LEBA to a diverse, multinational dataset^35^ to test whether light exposure-related behaviors predict self-reported sleep timing, sleep disturbances, and sleep-related impairment. We hypothesized that more outdoor time, stronger engagement in morning and daytime lighting practices, and greater control of evening light would be associated with earlier sleep timing and fewer sleep complaints, whereas more frequent device use in bed would relate to later sleep timing and more sleep complaints.

## Methods

### Participants

In total, 826 complete responses were obtained from an international, ageLdiverse sample. To ensure data quality, only participants who correctly answered all five embedded attention checks were included in the final sample, which comprised 774 individuals (383 women, 376 men, 15 other). Ages ranged from 11 to 84 years (*M* = 32.6 ± 14.6 y). To reach a broad and diverse population, the survey was made openly accessible worldwide and shared through multiple online channels. Specifically, participants were recruited via the website of the science communication project “Enlighten your clock” (Weinzaepflen & Spitschan (2021); https://enlightenyourclock.org/participate-in-research), which was co-released with this survey, as well as through social media platforms (LinkedIn, Twitter, Facebook), mailing lists, word of mouth, and the investigators’ professional networks. In addition, the survey link was shared with support from f.lux (f.lux Software LLC.). Eligibility criteria were intentionally broad to maximize generalizability. Participants had to be at least 11 years old and provide informed consent (or legal guardian consent for minors), and not have participated previously. No further exclusion criteria were applied.

### Measures

From the survey data, we analyzed self-reported sleep outcomes, sleep timing, sleep disturbances, sleep-related impairment, in relation to every light exposure behavior. In addition, participants completed questionnaires assessing chronotype, sleep environment, perceived light sensitivity, and, for underage participants, pubertal development stage. Demographic information considered in the analyses were age, sex, occupational status (work, school, other), and occupational setting (e.g., home office, face to face, etc.). All self-reports referred to the past four weeks, and scoring followed published instructions for each instrument.

### Light Exposure Behavior Assessment (LEBA)

Light exposure-related behaviors were measured using the long form of the LEBA questionnaire^35^, which includes 23 items rated on a five-point Likert scale (1 =”never” to 5 = “always”, with some items reverse coded). The instrument captures five subscales reflecting distinct behavioral patterns. Wearing blue-light filters (F1) assesses the habit of using blue-filtering, or orange-/red-tinted glasses during the day, either indoors or outdoors, as well as within one hour of attempting to fall asleep. Time spent outdoors (F2) reflects the amount of daily outdoor exposure, ranging from brief periods under 30 minutes to more than three hours as well as the habit of going outdoors within two hours after waking. Device use in bed (F3) assesses engagement with phones or smartwatches around sleep, including use within an hour before attempted sleep time, during nocturnal awakenings, and immediately after waking. Using light before bedtime (F4) captures behaviors to reduce evening artificial light exposure, including dimming phone and computer screens, enabling night-shift modes, and minimizing light when getting up during the night. Engagement in morning and daytime lighting practices (F5) encompasses behaviors such as using tunable or LED lighting to create a healthy light environment, using a desk lamp for focused work, employing a dawn-simulation alarm, and turning on lights immediately upon waking. Higher sum scores indicate more frequent engagement in the respective behavior. Items in F4 were reverse coded so that higher values represent greater restriction of evening light. Therefore, in the following, we refer to this factor as controlling light before bedtime. The LEBA questionnaire is available on Github (https://github.com/leba-instrument/leba-instrument-en/) as well as on the dedicated website of the LEBA inventory (https://leba-instrument.org/) under an open-access license (Creative Commons CC-BY).

### Sleep Timing

Sleep timing was assessed with the Munich Chronotype Questionnaire (MCTQ)^23^, using the adult or children’s version depending on participant age. From the MCTQ, we obtained mid-sleep on free days (MSF), defined as the midpoint between sleep onset and offset on days without social obligations (i.e., free days). We also calculated mid-sleep corrected for oversleep (MSFsc), which adjusts MSF for the sleep debt accumulated over the work week that is typically compensated by sleeping longer on free days^39^. The MCTQ also provided estimates of average weekly outdoor light exposure duration.

### Sleep Disturbance and Sleep-Related Impairment

Sleep problems were assessed using the Patient-Reported Outcomes Measurement Information System (PROMIS) Sleep Disturbance 4a and Sleep-Related Impairment short forms. PROMIS is a standardized system of patient-reported outcome measures built from large item banks developed using Item Response Theory, which allow brief, calibrated subsets of items to reliably assess the same underlying constructs^40^. The 4a forms used here consist of four items selected and calibrated from the adult item banks^41^ and, for participants completing the pediatric questionnaire, the corresponding pediatric short forms^42^. Items are rated on a five-point Likert scale (1 = “never” to 5 = “always”). Higher scores reflect more frequent sleep disturbances or sleep-related impairment.

### Light Sensitivity

Perceived sensitivity to light was assessed with the Photosensitivity Assessment Questionnaire (PAQ)^43^. The PAQ consists of 16 binary items (0 = “no”, 1 = “yes”), grouped into subscales for photophobia and photophilia. Higher subscale scores indicate greater light sensitivity or preference.

### Sleep Environment

Participants’ sleep environment was captured using the Assessment of the Sleep Environment Questionnaire (ASE)^44^. The ASE includes 13 items rated on a four-point Likert scale (0 = “strongly disagree” to 3 = “strongly agree”). Higher sum scores denote a more appropriate and supportive sleep environment.

### Pubertal Development

For participants younger than 18 years, pubertal stage was assessed with the Self-Rating Scale for Pubertal Development^45^. This scale comprises five items rated on a four-point Likert scale (1 = “not yet started” to 4 = “seems complete”), plus an additional item asking girls whether they had experienced menarche. Higher sum scores indicate more advanced pubertal development.

### Procedure

The anonymous online survey was administered via REDCap electronic data capture tools^46,47^ hosted at the University of Basel’s sciCORE facility. After providing informed consent, participants completed the online questionnaires. Completion took approximately 20 minutes and was not compensated. The study followed a quantitative, crossLsectional design, and the survey was openly accessible worldwide from 17 May 2021 until 24 November 2024, when data collection was closed prior to preregistering the analyses. To ensure data quality, four attention-check items were embedded throughout, with the following phrasing: “We want to make sure you are paying attention. What is 4+5?”; “Please select ‘Strongly disagree’ here.”; “Please type in ‘nineteen’ as a number.”; and “Please select ‘Does not apply/I don’t know.’ here.”.

Although the research project used fully anonymous data and therefore fell outside the scope of the Human Research Act, the cantonal Ethics Commission (Ethikkommission Nordwest-und Zentralschweiz, EKNZ) reviewed the project (ID Req-2021-00488) and issued an official clarification of responsibility. All procedures complied with the Declaration of Helsinki.

### Data analysis

All analyses were pre-registered (OSF: https://doi.org/10.17605/OSF.IO/SK9J2). The analytic approach comprised data preprocessing, confirmatory Bayesian modelling of preregistered hypotheses, and exploratory analyses to characterize broader associations among study variables.

During preprocessing, we noted that entries for bedtime and sleeptime in the MCTQ were especially prone to formatting errors or conceptual misunderstandings. To address this, entries given in a 12h format that plausibly reflected nocturnal sleep (defined as 12:00-06:00) were converted into 24h format. When bedtime and sleeptime were reversed but differed by no more than two hours (e.g., 23:00 and 21:30), values were swapped to restore plausibility, as sleeptime should follow bedtime. Remaining implausible responses were flagged and set to missing. The resulting amount of missing data was minimal and assumed to be random.

We tested the preregistered hypotheses using Bayesian linear models (lmBF function). For each sleep outcome (MSF and MSFsc, PROMIS sleep disturbances and sleep-related impairment), we compared a null model including age, sex, and work environment with four separate alternative models, each including one LEBA factor (for factors F2-F5). Evidence for each behavioral predictor was quantified with Bayes Factors (BFLL), indicating how much more likely the data were under the alternative model relative to the null. Values of BFLL > 1 indicate evidence in favor of the alternative, values < 1 indicate evidence for the null. Interpretation followed the Jeffreys’ scale adapted by Lee and Wagenmakers (2014), with BFLL ≥ 10 considered strong evidence for the alternative, BFLL ≤ 0.1 considered strong evidence for the null, and intermediate values reported as graded evidence^48^ (see Table 1). Bayes factors exceeding 1000 were reported as BFLL > 1000. To characterize effect estimates, we sampled 10,000 iterations from the posterior distribution of each model and extracted posterior means, standard deviations, and 95% credible intervals (CI). A credible interval reflects the range within which 95% of the posterior probability lies, given the data and the prior, there is a 95% probability that the true effect lies within this interval. If this interval excluded zero, we interpreted the effect as credibly different from zero. All models used the default priors implemented in the *BayesFactor* package.

**Table 1.** Classification Scheme for the Interpretation of Bayes Factors.

| BF $\square\square$ | Grade of evidence for the alternative hypothesis |
| --- | --- |
| 1-3 | Anecdotal evidence |
| 3-10 | Moderate evidence |
| 10-30 | Strong evidence |
| $\geq 30$ | Very strong evidence |

*Note.* BF $\square\square$ denotes the Bayes factor comparing the alternative hypothesis ( $H\square$ ) to the null hypothesis. Classification of Bayes factors follows the Jeffreys' (1961) evidence scale, as adapted by Lee and Wagenmakers (2014), for interpreting evidence in favor of $H\square$ <sup>48</sup>. BF $\square\square \geq 10$ were considered conclusive evidence for an effect. Abbreviations: BF = Bayes factor, $H\square$ = alternative hypothesis.

Exploratory analyses examined additional variables of interest beyond the preregistered hypotheses, including photophilia, photophobia, appropriateness of the sleep environment, sleep complaints, and weekly light exposure duration. Pairwise associations among numerical variables were evaluated using Spearman correlations (corr.test() function with pairwise deletion and false-discovery-rate correction). To identify broader multivariate patterns, we conducted a principal component analysis using Spearman correlation matrices. Sampling adequacy was assessed using the Kaiser-Meyer-Olkin measure (KMO)^49^ and Bartlett’s test of sphericity^50^. The number of components to retain was evaluated using parallel analysis, the minimum average partial criterion (MAP)^51^, and inspection of scree plots. Components were extracted using principal components and rotated using an oblique (oblimin) rotation to allow for correlated dimensions. Loadings were interpreted using a conservative cutoff of |0.40|. These exploratory analyses were not part of the confirmatory hypothesis testing and are presented in full in the Supplementary Material.

All analyses were conducted using R 4.4.2^52^ Bayesian models were estimated using the *BayesFactor* package^53^. MCTQ variables were computed with the *mctq* package^54^. Exploratory correlation analyses and PCA used the *psych* package^55^, and correlation visualizations were prepared with *ggcorrplot*^56^. The survey dataset and analysis code are available on Github (https://github.com/rrlazar/Sleep_And_Light_Exposure_Behaviour). The repository will be made publicly accessible upon publication.

## Results

Nearly half of the sample (365 participants, 47%) reported English as their native language. Participants were located across 75 countries and 28 time zones, with the largest proportions residing in London, United Kingdom (n = 86, 11.1%), New York, United States (n = 68, 8.8%), and Berlin, Germany (n = 64, 8.3%). Occupational status was diverse, with 438 participants (57%) employed, 211 (27%) students, and 125 (16%) neither employed nor studying. Work settings still reflected the impact of the COVID-19 pandemic, as 322 (42%) reported working remotely, and 169 (22%) working in hybrid settings. Descriptive statistics of the main study variables are presented in Table 2. For the LEBA subscales, score ranges differed according to the number of items contributing to each factor. Higher sum scores indicated more frequent engagement in the respective behavior, with minimal scores reflecting that the behavior is never exhibited and maximal scores indicating it is always exhibited. Across factors, mean values fell near the midpoints of their possible ranges, suggesting that participants engaged in the assessed behaviors to varying degrees. Sleep outcomes showed intermediate levels of disturbance and impairment. For both PROMIS subscales, which range from 4 (never) to 20 (always), participants reported mean scores of 10.8 (±3.6) for sleep disturbances and 10.1 (±4.2) for sleep-related impairment. Midsleep time also fell largely within the intermediate chronotype range. Participants showed an average MSF of 04:52 (± 02:09) and an average MSFsc of 04:28 (± 02:15). According to the chronotype categories suggested by Zhang et al (2019)^57^, based on cut-offs introduced by Roennberg (2007)^39^, these values correspond to an intermediate chronotype (<3:00 early type, 3:00-5:00 intermediate, >5:00 late type). Weekly outdoor light exposure derived from the MCTQ was low overall (*M* = 1.9h per week), aligning with prior findings that modern populations receive limited daylight. Additional descriptive statistics for the light sensitivity scale (PAQ), sleep environment (ASE), and pubertal development scale (PDS) are also shown in Table 2.

**Table 2.**
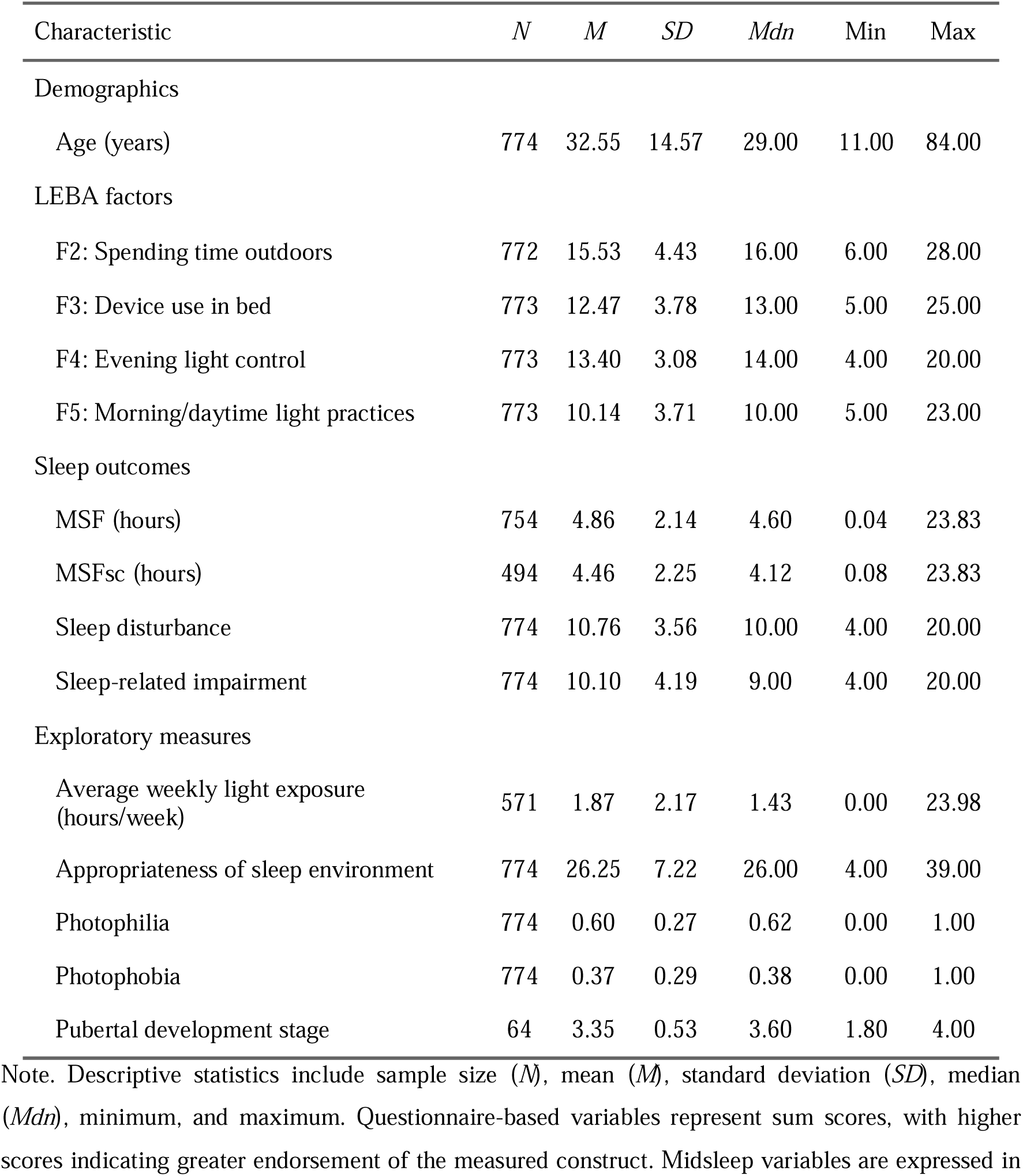
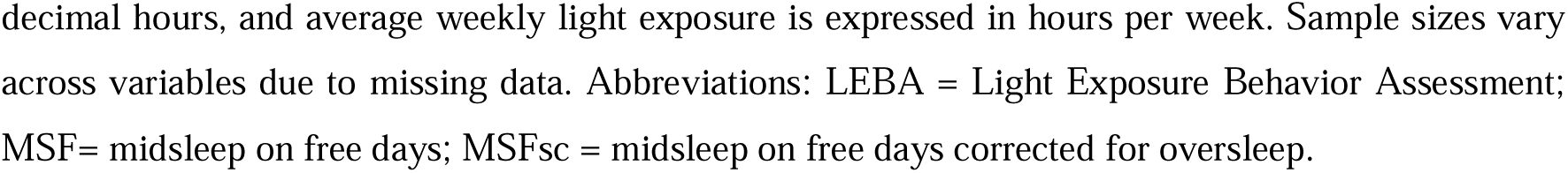
Descriptive Characteristics of Demographic, Light Exposure, and Sleep-Related Variables.

| Characteristic | *N* | *M* | *SD* | *Mdn* | Min | Max |
| --- | --- | --- | --- | --- | --- | --- |
| Demographics |  |  |  |  |  |  |
| Age (years) | 774 | 32.55 | 14.57 | 29.00 | 11.00 | 84.00 |
| LEBA factors |  |  |  |  |  |  |
| F2: Spending time outdoors | 772 | 15.53 | 4.43 | 16.00 | 6.00 | 28.00 |
| F3: Device use in bed | 773 | 12.47 | 3.78 | 13.00 | 5.00 | 25.00 |
| F4: Evening light control | 773 | 13.40 | 3.08 | 14.00 | 4.00 | 20.00 |
| F5: Morning/daytime light practices | 773 | 10.14 | 3.71 | 10.00 | 5.00 | 23.00 |
| Sleep outcomes |  |  |  |  |  |  |
| MSF (hours) | 754 | 4.86 | 2.14 | 4.60 | 0.04 | 23.83 |
| MSFsc (hours) | 494 | 4.46 | 2.25 | 4.12 | 0.08 | 23.83 |
| Sleep disturbance | 774 | 10.76 | 3.56 | 10.00 | 4.00 | 20.00 |
| Sleep-related impairment | 774 | 10.10 | 4.19 | 9.00 | 4.00 | 20.00 |
| Exploratory measures |  |  |  |  |  |  |
| Average weekly light exposure (hours/week) | 571 | 1.87 | 2.17 | 1.43 | 0.00 | 23.98 |
| Appropriateness of sleep environment | 774 | 26.25 | 7.22 | 26.00 | 4.00 | 39.00 |
| Photophilia | 774 | 0.60 | 0.27 | 0.62 | 0.00 | 1.00 |
| Photophobia | 774 | 0.37 | 0.29 | 0.38 | 0.00 | 1.00 |
| Pubertal development stage | 64 | 3.35 | 0.53 | 3.60 | 1.80 | 4.00 |

Note. Descriptive statistics include sample size (*N*), mean (*M*), standard deviation (*SD*), median (*Mdn*), minimum, and maximum. Questionnaire-based variables represent sum scores, with higher scores indicating greater endorsement of the measured construct. Midsleep variables are expressed in

### Confirmatory Analyses

#### Sleep Timing

We first examined whether light exposure-related behaviors predicted sleep timing, using MSF (*n* = 751) and MSFsc (*n* = 491). The smaller sample for MSFsc reflects the MCTQ requirement that oversleep corrected values are only valid when no alarm is used on free days^58^. Overall, our results strongly support the pre-registered hypothesis that more outdoor time is associated with earlier sleep timing. For both MSF and MSFsc, models including outdoor time showed extreme evidence in favor of an effect (BFLL > 1000), and posterior estimates (see Supplements) consistently indicated earlier mid-sleep among individuals who spent more time outdoors. For the remaining behaviors, however, the pre-registered hypotheses were not supported. For MSF, the models including morning and daytime lighting practices (BFLL = 0.42), device use in bed (BFLL = 0.19), and control of evening light (BFLL = 0.35) provided insufficient evidence for an association. Results for MSFsc showed the same pattern. Again, morning and daytime light practices (BFLL = 0.19), device use in bed (BFLL = 0.15), and control of evening light (BFLL = 0.15) yielded insufficient evidence, and none of the corresponding posterior estimates were credibly distinct from zero. Full posterior summaries are reported in the Supplemental Material.

#### Sleep Disturbances

Next, we tested whether LEBA factors predicted sleep disturbances (*n* = 771). Overall, the results provide strong and consistent evidence that time spent outdoors related to fewer sleep disturbances, and that greater device use in bed related to more disturbances, whereas the expected beneficial effects of morning and daytime lighting practices and evening light control were not supported. The model including outdoor time showed extreme evidence in favor of an effect (BFLL > 1000), with posterior estimates indicating that individuals who spent more time outdoors reported fewer sleep disturbances. Device use in bed likewise showed extreme evidence in favor of H1 (BFLL > 1000), with posterior estimates indicating that greater device use in bed is associated with more sleep disturbances. For the two remaining factors, morning and daytime lighting practices (BFLL = 0.24) and evening light control (BFLL = 0.31) yielded moderate evidence against an effect, yet the credible intervals were not credibly distinct from zero. These results indicate that sleep disturbances were reliably and meaningfully related only to outdoor light exposure and device use in bed.

#### Sleep-Related Impairment

Finally, we examined whether LEBA factors predicted sleep-related impairment (*n* = 771). Overall, the results closely resembled the findings for sleep disturbances, with strong and reliable associations for outdoor time and device use in bed, whereas the expected effects of morning and daytime lighting practices and evening light control were not supported. As with the other outcomes, spending time outdoors showed extreme evidence in favor of H1 (BFLL > 1000), with posterior estimates indicating lower sleep-related impairment in individuals with higher time spent outdoors. Device use in bed also showed extreme evidence in favor of HL (BFLL > 1000), with posterior estimates suggesting that more frequent device use in bed was associated with greater impairment. For the remaining behaviors, the findings were not confirmatory. Morning and daytime lighting practices (BFLL = 1.81) and evening light control (BFLL = 1.40) provided insufficient evidence. Although their posterior estimates suggested small, positive associations, these behaviors did not meaningfully improve model fit and should not be further interpreted as robust effects. Thus, only outdoor time and device use in bed showed reliable associations with sleep-related impairment, whereas the effects of morning and daytime lighting practices and evening light control remain inconclusive.

**Figure 1.**
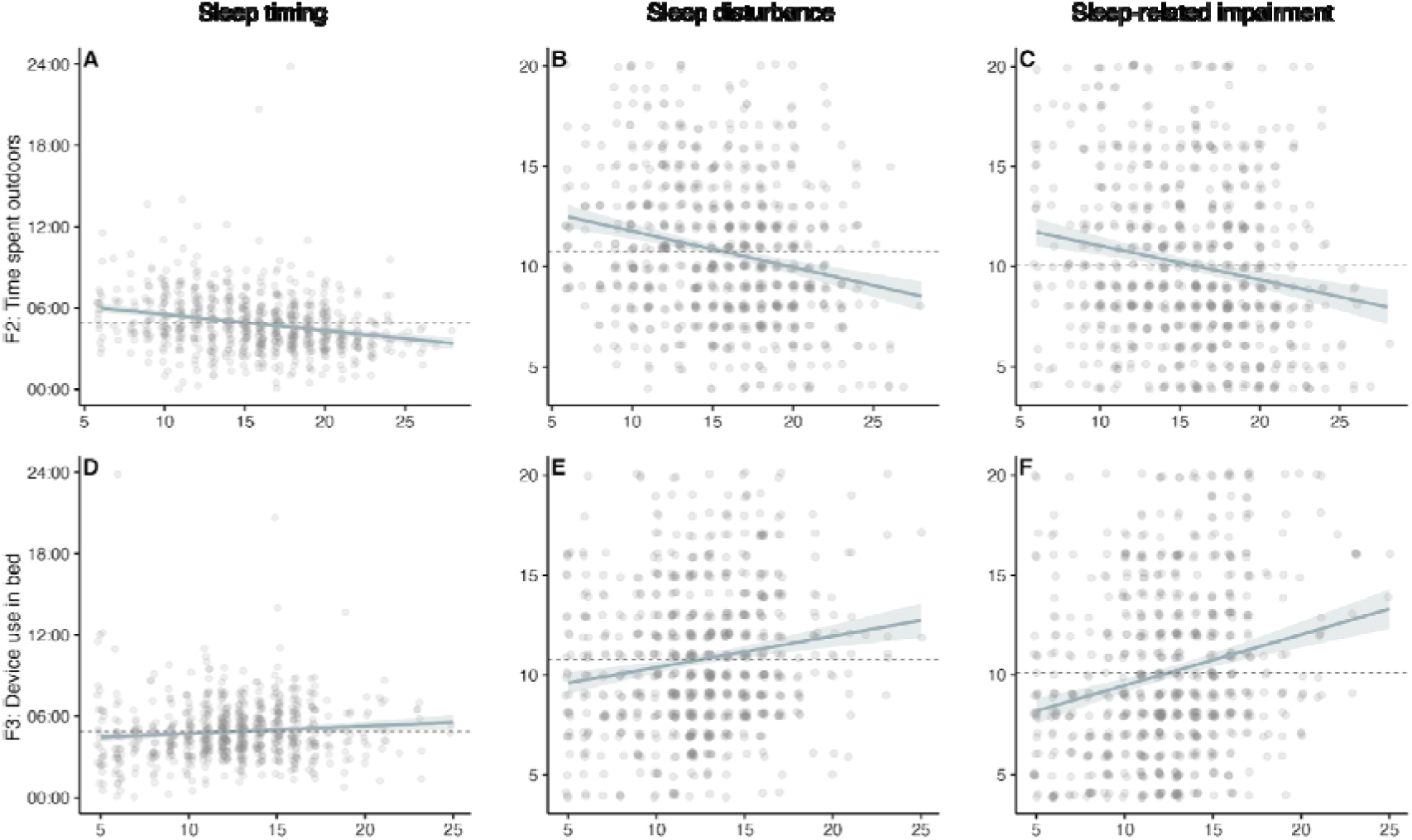
Differential Associations of Time Outdoors and Device Use in Bed With Sleep Outcomes *Note.* Scatterplots show associations between LEBA factor sum scores and sleep-related outcomes. The top row (Panels A-C) shows associations for time spent outdoors (F2), and the bottom row (Panels D-F) shows associations for device use in bed (F3). Columns represent sleep timing (midsleep on free days), sleep disturbance, and sleep-related impairment, respectively. Points represent individual participants and are jittered horizontally to reduce overplotting. Solid lines indicate linear regression fits, with shaded bands representing 95% confidence intervals. Dashed horizontal lines denote the sample mean of the respective outcome. Sleep timing is expressed as clock time (hh:mm), whereas sleep disturbance and sleep-related impairment represent questionnaire sum scores, with higher values indicating more occurrence. More frequent time spent outdoors was associated with earlier sleep timing and fewer sleep-related concerns. In contrast, more frequent device use in bed was associated with more sleep-related concerns, albeit without conclusive evidence regarding sleep timing. Analyses were based on data from *N* = 751 individuals for midsleep on free days (Panels A and D), and *N* = 771 individuals for sleep disturbance and sleep-related impairment due to missing data (Panels B, C, E, and F).

## Discussion

In this preregistered study, we found that among the light exposure-related behaviors assessed, time spent outdoors and device use in bed were the most meaningful behaviors linked to self-reported sleep outcomes. Time spent outdoors, reflecting the amount of natural daylight individuals were exposed to daily, showed the clearest and most robust associations: There was extreme evidence that individuals who spent more time outdoors went to sleep earlier, experienced fewer sleep disturbances, and reported less sleep-related daytime impairment. In contrast, more frequent device use in bed, capturing the tendency to use phones or smartwatches around the sleep period, was reliably associated with poorer sleep, particularly more disturbances during the night and greater impairment the next day. By comparison, morning and daytime lighting practices, describing how individuals shape their indoor light environment after waking and across the day, and evening light control, referring to efforts to reduce evening and night-time light exposure, were not meaningfully related to sleep timing and disturbances, and showed only weak, anecdotal associations with sleep-related impairment.

Time spent outdoors consistently emerged as the light exposure-related behavior most relevant to the sleep outcomes assessed, with individuals who spent more time outdoors reporting earlier sleep timing, fewer sleep disturbances, and less sleep-related daytime impairment. This beneficial pattern aligns well with a large body of evidence showing that light influences sleep through both circadian and homeostatic processes. The ipRGC system is driven by light input, and natural daylight provides a particularly effective stimulus^59^. Daylight exceeds typical indoor illuminance by several orders of magnitude, often reaching 25,000 lux in full daylight and up to 100,000 lux in direct sun, compared with approximately 300 to 500 lux in office environments^60,61^. While bright light in the morning advances circadian phase and delays it in the evening, bright light, particularly light rich in short-wavelengths, also elicits acute alerting effects^18,62^ that can influence sleep latency and daytime functioning^21,63^. Field-based studies similarly report that greater outdoor light exposure is associated with earlier sleep timing, better subjective sleep, and reduced daytime sleepiness^27,36,58^. More recent work suggests that morning sunlight may be particularly beneficial for advancing mid-sleep timing and improving perceived sleep quality^19^. Our findings add to this evidence by showing that a simple behavioral indicator of daylight exposure, self-reported daily time spent outdoors, corresponds not only to differences in sleep timing but also to sleep disturbances and daytime impairment in a large, diverse international sample. This is particularly relevant in modern environments, which are characterized by low daytime light intensities that contrast with the brighter days under which humans evolved. Because modern light exposure increasingly reflects behavioral choices, rather than environmental illumination alone^64^, our results highlight that engagement with outdoor daylight meaningfully relates to sleep outcomes and daily functioning. Exploratory associations further supported this behavioral interpretation, as individuals who spent more time outside also reported higher photophilia, lower photophobia, and substantially greater weekly light exposure duration. Therefore, our findings reinforce the well-established biological importance of daylight and additionally suggest that behavior-driven differences in daylight exposure carry measurable consequences for sleep in everyday life.

Using electronic devices in bed emerged as the second most influential behavior in our analysis showing reliable associations with more sleep disturbances and greater sleep-related daytime impairment. This is in line with evidence that evening screen exposure can exert both circadian and alerting effects. Light emitted from digital devices, which is typically enriched in short-wavelengths, has been shown to suppress the evening rise in melatonin^28,29^ and to delay circadian phase when exposure occurs before sleep^65^. Screens can also exhibit acute alerting effects^28^, which often accompany difficulties initiating sleep and poorer sleep continuity^66,67^. Accordingly, numerous population-based studies report that bedtime screen use is associated with later sleep timing, poorer sleep quality, greater daytime sleepiness^36,68,69^. More recent work, however, suggests that the downstream effects of evening light exposure on sleep may be less pronounced than previously assumed. Although evening screen light reliably suppresses melatonin, some studies did not observe corresponding alterations in sleep timing or structure once the light stimulus was removed^31^. Moreover, prior bright light exposure appears to mitigate even melatonin suppression and effects on sleep parameters^30,70^. These findings, together with proposals that mechanisms other than bright light, such as arousal, night-time disruption, or sleep displacement, may contribute to technology-related sleep complaints, have complicated the interpretation of screen-use effects. Indeed, most studies that measured objective sleep parameters report relatively small effect sizes, raising questions about their practical significance^32^. Our findings fit within this more mixed evidence base. Individuals reporting more frequent device use in bed also reported more sleep disturbances and greater daytime impairment, consistent with studies identifying adverse consequences of screen use near bedtime. At the same time, we did not observe an association with sleep timing. Although the Bayes factor provided moderate support for including device use in the sleep-timing model, the corresponding credible interval indicated that any true effect is likely small. This may reflect the limited impact of real-world device use, or the fact that LEBA items assess device use in bed but not duration, screen brightness, or blue-light-filter settings, factors known to modulate physiological responses^18,71^. It is also possible that directionality differs from a simple causal pathway, such that individuals experiencing more sleep problems use devices more as a coping or distraction strategy^32^. Taken together, our results support an association between device use in bed and subjective sleep complaints, while suggesting that its influence on sleep timing is limited, context-dependent, or requires finer-grained assessment to detect reliably.

Evening light control yielded inconclusive evidence with our sleep outcomes. Several factors may explain this limited pattern. The LEBA items capture behaviors such as dimming screens, enabling night shift mode, or reducing light at night, but they do not quantify parameters that determine physiological impact, such as illuminance, spectral content, exposure duration, or direction of light. Because circadian and alerting responses depend on retinal irradiance^34^, self-reported behaviors may not correspond to meaningful physiological differences. Although laboratory studies demonstrate sufficiently bright evening light can delay circadian timing and impair sleep^e.g., 28,66,72^, these effects typically rely on controlled and prolonged exposures. By contrast, real-world evening light is often shorter, more variable, and substantially less intense. For instance, LED and e-reader light can disrupt sleep in controlled studies^73,74^, but the exposure durations may not reflect habitual real-world screen use. Moreover, dimming a smartphone or activating night-shift mode considerably reduces melanopic activation^60^, making measurable effects less likely. Notably, even these widely recommended strategies do not consistently improve sleep. Duraccio and colleagues (2021) found that enabling night-shift mode did not reliably enhance sleep and, under some circumstances, was associated with more wakefulness, whereas abstaining from phone use yielded the best outcomes^75^. Evening light effects also depend on factors not captured by the LEBA, such as prior light exposure, ambient indoor lighting, and light exposure across a longer pre-sleep window^61,76^. Consistent with our findings, Siraji et al. (2023) also reported no meaningful associations for this behavioral domain^36^. Taken together, these results suggest that the LEBA operationalization of evening light control may not sufficiently reflect variation in physiologically relevant evening light exposure.

A comparable pattern emerged for behaviors shaping morning and daytime lighting practices, for which we likewise did not observe conclusive evidence for or against an effect on the sleep outcomes assessed. Although morning and daytime light can benefit circadian entrainment and sleep when delivered at sufficient intensity and spectral power^60,71^, the LEBA items in this domain, that is, questions relating to increasing illumination after waking and using adjustable, task-oriented lighting across the day, are only loosely linked to the retinal light dose that drives circadian physiology. Typical indoor lighting provides relatively low illuminance and limited short-wavelength content compared to outdoor light^61^, and self-reported behaviors cannot capture the substantial variability introduced by lighting geometry, device settings, or surrounding environmental context. Individuals scoring higher on this factor tended to report greater photophilia, suggesting that these practices may reflect preferences for brighter or more adapted lighting environments rather than behaviors that meaningfully alter physiological light exposure. Such preference-driven habits may therefore not translate into strong or effects on sleep in a real-world context. Cultural and geographical variation in daily light routines, including daylight availability and lighting preferences, may further contribute to inconsistent associations in our international sample^64^. Although Siraji and colleagues (2023) found that engagement in these morning and daytime light exposure-related behaviors predicted earlier chronotype and better perceived sleep quality in a Malaysian sample^36^, we did not replicate these associations in our larger and more heterogeneous sample. Overall, the inconclusive effects in our data suggest that these behaviors may play a less important role for sleep timing, sleep disturbances, and sleep-related impairment under everyday conditions, or that their impact is more difficult to capture with this behavioral self-report.

Several strengths characterize this work, alongside limitations inherent to its design. Applying the validated Light Exposure Behavior Assessment (LEBA) questionnaire to a large international sample (>700 participants across multiple time zones and age groups), we linked multiple light exposure-related behaviors to sleep outcomes under real-world conditions. Most studies so far have focused on the physical measurement of light exposure; wearable sensors and optical radiation dosimeters now allow continuous, real-world monitoring of light intensity and spectrum, providing ecologically valid estimates of retinal exposure^77–79^. However, while such methods can capture environmental exposure, they are intrusive, require continuous engagement, and vary in spectral sensitivity, placement, and adherence^80,81^. Moreover, given our modern society, light exposure results as a combination of environmental conditions and behavioral choices to shape individual light environments. While the LEBA cannot account for all environmental factors influencing light exposure, focusing on behavior complements sensor-based approaches by addressing volitional components of light use. Finally, online, voluntary recruitment may overrepresent younger or health-interested individuals, yet the geographic and demographic diversity of the sample enhances the robustness and ecological relevance of the findings. Although the cross-sectional design precludes causal inference, the observed associations offer valuable insights for applications and future research. Such future research should clarify how self-reported light behaviors translate into physiological light exposure. Combining the LEBA with objective field measurements would help determine how well behavioral tendencies map onto actual retinal illuminance. Studies spanning different seasons, latitudes, and cultural contexts could also test the generalizability of these findings and identify subgroups for whom light-related behaviors matter more or less. Longitudinal and within-person designs may further reveal information about the directionality and causality of effects and whether the small effects observed for morning and daytime lighting practices and evening light behaviors accumulate across time or emerge only under specific environmental conditions.

In conclusion, our findings suggest that everyday light exposure-related behaviors, particularly time spent outdoors and using electronic devices, are meaningfully associated with interindividual differences in sleep. Viewing light exposure as a modifiable behavior rather than only as an environmental condition offers a valuable perspective on how daily routines shape sleep in real-world settings. By applying this behavioral framework in a large and diverse sample, we demonstrate the utility of assessing light-related behaviors with the LEBA and highlight the potential for such tools to complement physiological and sensor-based approaches. These results support the broader view that healthier sleep emerges from an interplay between environmental conditions and individual choices. They also support recent proposals to consider light exposure as part of a healthy spectral diet throughout the day in both the general population and clinical settings. Evidence suggests that brighter days and darker evenings support circadian alignment, mood, and well-being, whereas the opposite can have adverse effects^e.g., 82^. Our findings extend this perspective by showing that time spent outdoors is also reliably associated with sleep outcomes in everyday life. Importantly, treating light exposure as a health behavior highlights what individuals can actively do to support their sleep. Small, accessible behaviors may help create conditions that promote better sleep in real-world settings. Such behavioral adjustments contribute to what has been defined as sleep health, that is, the adaptation of the sleep-wake pattern to individual, social, and environmental demands in a way that supports physical and mental well-being^83^. Together, our findings underscore sleep health as both an individual and societal priority and motivate further research into the environmental and behavioral factors that determine healthy sleep.

## Author contributions

Conceptualization and data curation: RL and MS. Formal analysis: ASL and RL. Supervision: CB. Visualization: ASL. Writing – original draft: ASL. Writing – review and editing: ASL, RL, MS, and CB.

## Data availability

The LEBA questionnaire is available on Github (https://github.com/leba-instrument/leba-instrument-en/). The survey dataset and analysis code are available on Github (https://github.com/rrlazar/Sleep_And_Light_Exposure_Behaviour) and will be made publicly accessible upon publication.

## Declaration of generative AI and AI-assisted technologies in the writing process

During the preparation of this manuscript, the authors used ChatGPT (OpenAI, chatgpt.com) to support coding tasks (i.e., troubleshooting and plotting) and for language polishing, including improving clarity, coherence, and wording. All outputs were critically reviewed and edited by the authors, who take full responsibility for the content of the publication.

