## Supplementary Material for "Associations Between Habitual Light Exposure-Related Behaviors and Sleep Timing and Sleep Complaints in an International Community Sample"

### Supplementary Results

#### Suppl. Figure 1

*Posterior Effect Size Estimates for Time Spent Outdoors and Device Use in Bed in Relation to Sleep Outcomes*

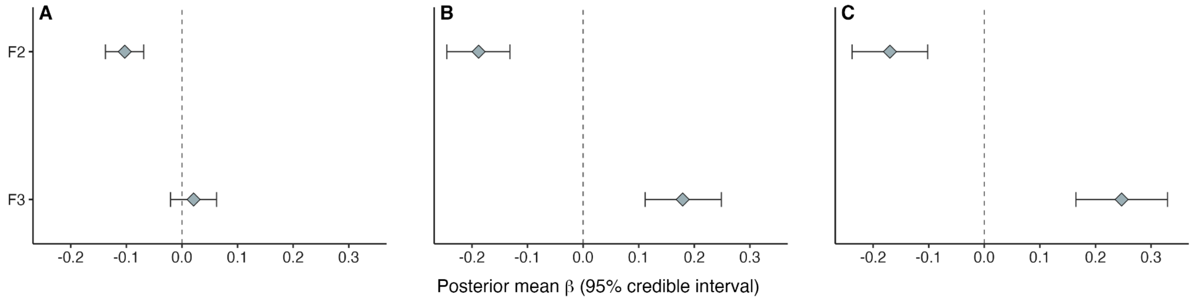


*Note.* This figure demonstrates posterior mean regression coefficients (β) and 95% credible intervals for associations between the most informative LEBA behaviors and sleep outcomes: sleep timing (A), sleep disturbances (B), and sleep-related impairment (C). Within each panel, estimates are shown for time spent outdoors (F2) and device use in bed (F3). Points indicate posterior mean effect size estimates, and horizontal lines denote 95% credible intervals. The dashed vertical line marks the null effect ($\beta = 0$). Estimates are derived from Bayesian regression models.

**Suppl. Table 1**

*Posterior Estimates for Midsleep on Free Days (MSF) in the Assessed LEBA Behavior Domains*

| Predictor | $BF_{10}$ | Estimate | SD | 95% CI (lower; upper) |
|---|---|---|---|---|
| F2: Time spent outdoors | >1000 | -0.10 | 0.02 | [-0.14; -0.07] |
| F3: Device use in bed | 0.19 | 0.02 | 0.02 | [-0.02; 0.06] |
| F4: Evening light control | 0.40 | 0.04 | 0.02 | [-0.01; 0.09] |
| F5: Morning and daytime lighting practices | 0.41 | -0.03 | 0.02 | [-0.07; 0.01] |

Abbreviations: $BF_{10}$ = Bayes factor for the alternative model over the null model; SD = standard deviation; CI = credible interval.

**Suppl. Table 2**

*Posterior Estimates for Midsleep on Free Days Corrected for Oversleep (MSFsc) in the Assessed LEBA Behavior Domains*

| Predictor | $BF_{10}$ | Estimate | SD | 95% CI (lower; upper) |
|---|---|---|---|---|
| F2: Time spent outdoors | >1000 | -0.10 | 0.02 | [-0.14; -0.06] |
| F3: Device use in bed | 0.16 | 0.01 | 0.03 | [-0.04; 0.06] |
| F4: Evening light control | 0.16 | 0.01 | 0.03 | [-0.06; 0.07] |
| F5: Morning and daytime lighting practices | 0.19 | 0.02 | 0.03 | [-0.03; 0.07] |

Abbreviations: $BF_{10}$ = Bayes factor for the alternative model over the null model; SD = standard deviation; CI = credible interval.

**Suppl. Table 3**

*Posterior Estimates for Sleep Disturbances in the Assessed LEBA Behavior Domains*

| Predictor | $BF_{10}$ | Estimate | SD | 95% CI (lower; upper) |
|---|---|---|---|---|
| F2: Time spent outdoors | >1000 | -0.19 | 0.03 | [-0.24; -0.13] |
| F3: Device use in bed | >1000 | 0.18 | 0.04 | [0.11; 0.25] |
| F4: Evening light control | 0.31 | 0.06 | 0.04 | [-0.03; 0.14] |
| F5: Morning and daytime lighting practices | 0.24 | -0.04 | 0.03 | [-0.11; 0.03] |

Abbreviations: $BF_{10}$ = Bayes factor for the alternative model over the null model; SD = standard deviation; CI = credible interval.

**Suppl. Table 4**

*Posterior Estimates for Sleep-Related Impairment in the Assessed LEBA Behavior Domains*

| Predictor | $BF_{10}$ | Estimate | SD | 95% CI (lower; upper) |
|---|---|---|---|---|
| F2: Time spent outdoors | >1000 | -0.17 | 0.03 | [-0.24; -0.10] |
| F3: Device use in bed | >1000 | 0.25 | 0.04 | [0.17; 0.32] |
| F4: Evening light control | 1.33 | 0.11 | 0.05 | [0.01; 0.20] |
| F5: Morning and daytime lighting practices | 1.58 | 0.09 | 0.04 | [0.01; 0.17] |

Abbreviations: $BF_{10}$ = Bayes factor for the alternative model over the null model; SD = standard deviation; CI = credible interval.

### Exploratory analyses

A correlation matrix of the numerical variables selected for further exploratory investigation is shown in Suppl. Figure 2. Overall, correlations were small to moderate in magnitude. Correlations among the LEBA factors were limited, with only weak associations observed between time spent outdoors (F2) and device use in bed (F3), and between F2 and engagement in morning and daytime lighting practices (F5; ρ = .13), suggesting that the LEBA factors capture largely independent behavioral tendencies. Several associations with external variables reached significance. Wearing blue light filters correlated positively with photophobia (ρ = .18). Time spent outdoors showed a consistent pattern of associations as seen in confirmatory analyses, correlating positively with weekly light exposure duration (ρ = .65) and photophilia (ρ = .29), but negatively with sleep timing (ρ = -.28 for MSF and ρ = -.3 for MSFsc), and photophobia (ρ = -.33). Device use in bed showed a small positive association with sleep disturbance (ρ = .17). Associations with sleep timing, sleep-related impairment, and other variables were weak (|ρ| ≤ .13). Evening light control showed no meaningful correlations with sleep-related outcomes, and engagement in morning and daytime lighting practices correlated positively with photophilia (ρ = .21).

**Suppl. Figure 2**

*Correlation Matrix for Exploratory Analysis of Additional Study Variables*

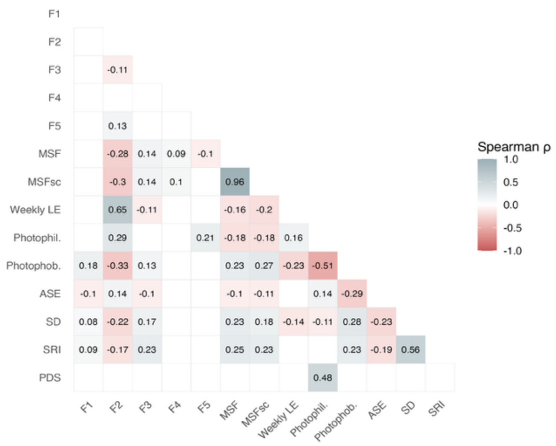


*Note.* Spearman rank-order correlations (ρ) were computed using pairwise-complete observations with false discovery rate (FDR) correction. Only significant correlations (p < .05, FDR-adjusted) are shown.

To identify latent structures beyond pairwise associations, we conducted a principal component analysis including the LEBA factors, MSF, average weekly light exposure duration, photophobia and photophilia scores, sleep environment appropriateness, and PROMIS sleep disturbance and sleep-related impairment. Sampling adequacy was acceptable (KMO = 0.68)[1], and Bartlett's test of sphericity[2] indicated that the correlation matrix was suitable for component analysis ($\chi^2(66) = 1091.98$, $p < .001$). The number of components to retain was evaluated using parallel analysis, the minimum average partial (MAP) criterion, and inspection of the scree plot. Parallel analysis suggested a two-component solution, whereas MAP indicated a single dominant component. Given the exploratory aim and interpretability of the solution, we retained two components and applied oblique rotation.

Using a conservative loading cutoff of |0.40|, Component 1 was characterized by high loadings of sleep disturbance, sleep-related impairment, and device use in bed (F3), reflecting a dimension of sleep complaints and associated device use consistent with the confirmatory results. Component 2 showed strong positive loadings of photophilia, spending time outdoors (F2), and engagement in morning and daytime lighting practices (F5), alongside a negative loading of photophobia, capturing a dimension of light affinity and active light exposure behaviors. Sleep timing and sleep environment appropriateness showed moderate loadings at a lower threshold but did not meet the predefined cutoff for interpretation. Together, the two components retained accounted for 36% of the total variance. Weekly light exposure introduced substantial missingness and reduced sample size (n = 556 vs. n = 751) without altering the substantive component structure. Therefore, we conducted the PCA without average weekly light exposure to increase interpretability.

**Suppl. Table 5**

*Rotated Component Loadings for the Two-Component PCA Solution*

| Variable | Component | |
|---|---|---|
| | Component 1 | Component 2 |
| F3: Device use in bed | .49 | |
| Sleep disturbances | .73 | |
| Sleep-related impairment | .81 | |
| F2: time spent outdoors | | .58 |
| F5: morning and daytime lighting practices | | .55 |
| Photophilia | | .80 |
| Photophobia | | -.67 |

*Note.* Loadings < .40 are not shown.
